# Deep Integrated Network Analysis – a data-driven tool to discover and characterize disease pathways in the liver

**DOI:** 10.1101/2025.03.17.643687

**Authors:** Jaclyn E. Quin, Miren Urrutia Iturritza, Katherine D. Mosquera, Franziska Hildebrandt, Fredrik Barrenäs, Johan Ankarklev

## Abstract

**Background & Aims:** An extensive number of studies have utilized transcriptomic profiling as a valuable tool for uncovering genes related to diseases and physiological processes of the liver. Here we combine this wealth of information to provide a powerful resource, by computationally constructing a comprehensive and unbiased network of gene interactions specific to the liver.

**Methods:** We have performed a computational approach termed Deep Integrated Network Analysis (DINA) on a curated catalog of 655 liver transcriptomic datasets (including a total of 48,311 transcriptomes). These datasets include human, monkey, mouse, rat and other mammalian species, and studies linked to a broad range of conditions. Together this facilitated construction of a network of strongly conserved gene-gene interactions relevant across the spectrum of liver diseases.

**Results:** The ‘Liver DINA Resource’ described herein contains 89,683 statistically conserved interactions among 19,317 genes in a unified network unique to the mammalian liver. The network unveils a hierarchical structure of strongly co-regulated modules, which are organized into a Tree-and-Leaf Network to provide a comprehensive overview of the resource.

**Conclusions:** This data-driven resource provides an interactive tool for the examination of previously undescribed gene networks, and enables unbiased analysis of transcriptomic datasets of the liver, thus preventing bias in favor of well-studied genes and pathways and providing a complementary approach towards novel discoveries.

## Introduction

The liver is a vital organ with multifaceted functions, processing the nutrient-rich blood leaving the gastrointestinal tract, regulating nutrient levels in the blood and preventing ingested harmful substances from entering circulation. A spectrum of liver diseases have a significant impact on human health, causing approximately 1 in 25 deaths worldwide ^1^. Its causes include infections, toxins, alcohol, metabolic-associated fatty liver disease (MAFLD), and others. Particularly, viral hepatitis B- and C-related diseases lead to over 1 million deaths each year ^1^. However, while the management of viral hepatitis is improving, the prevalence of MAFLD and alcohol-associated liver disease (ALD) is rising, especially in socio-economically developed regions of the world ^2^. Regardless of the cause, liver injury manifests as hepatitis; chronic hepatitis progresses through fibrosis to irreversible cirrhosis and ultimately leads to chronic liver failure. All these common causes of chronic hepatitis are also risk factors for hepatocellular carcinoma (HCC), one of the most common malignant tumors in the world ^3^. Together, these global health developments underscore the importance of improving our understanding of liver diseases and their causes.

The liver has been studied extensively using transcriptomic approaches, which have provided valuable insights to the field, and are becoming increasingly sophisticated, for example incorporating spatial and/or single-cell sequencing of the liver in obesity ^4^, ALD ^5^, infection ^6–8^, steatosis ^9,10^, fibrosis ^11^, and tissue repair ^12^. Collectively, the wealth of publicly available liver transcriptomic datasets creates a powerful opportunity to objectively investigate liver physiology and disease mechanisms, without relying on previously known gene functions and cellular pathways.

Integrating transcriptomic data has previously been used to create bioinformatic resources, both universal, e.g. GeneMania ^13^, STRING ^14,15^, FunCoup ^16^, as well as specific for a limited number of tissues, e.g. blood ^17^. Here, we compile an extensive catalog of transcriptomic datasets, specifically for liver disease studies. To this end, we incorporate multiple species and a broad range of liver conditions. This strategy, termed Deep Integrated Network Analysis (DINA), entails a computational ‘guilt-by-association’ approach to both identify, and statistically annotate, preserved gene-gene interactions across the entire dataset catalog. This enables us to construct a comprehensive network that encompasses shared and unique pathways across the spectrum of liver conditions. As our approach is fully data-driven and not biased by the extent of prior knowledge, this resource is inherently suited for novel discoveries ^18^.

## Materials and methods

### Dataset inclusion criteria

Publicly available liver datasets were accessed from the Gene Expression Omnibus of the National Center for Biotechnology Information (NCBI GEO) ^19,20^. Inclusion criteria were: (i) study type was either ‘expression profiling by array’ or ‘expression profiling by high throughput sequencing’; (ii) taxonomic group was ‘mammals’; (iii) the dataset included ≥ 5000 human genes, or one-to-one human gene orthologs; and (iv) sample size ≥ 10. GEO Series which included two or more species were separated into one dataset for each species.

### Data processing and filtering

Raw read counts from each RNA sequencing study were obtained as uniformly processed raw read counts from ArchS4 ^21^, where possible. The remaining RNA sequencing or microarray datasets were obtained from GEO. All gene or probe IDs were annotated to human Ensembl gene IDs, with non-human genes converted to human one-to-one gene orthologues ^22^.

### Quality assessment for inclusion of datasets

Each dataset first underwent rank-normalization followed by gene-gene correlation using biweight midcorrelation, and finally complete linkage clustering. Each dataset’s dendrogram was cut at a height that produced 50 gene clusters. Pairwise similarity between cluster membership vectors was measured using normalized mutual information (NMI). Each NMI value was calculated as a z-score, with the random distribution for NMI values calculated using a permutation test where one cluster membership vector was randomized 1,000 times. The similarity between the datasets was then analyzed using a k-nearest neighbor network (KNN), connecting each dataset to four other datasets based on highest z-score (k=4).

### Network inference

The gene-to-gene network was obtained through the Mavatar Discovery platform (Mavatar Ltd., Sweden) during alpha testing in Q4 2024. Data processing and filtering, quality assessment for inclusion of datasets, and network inference are part of the platform’s pipeline. Briefly, the platform identifies gene pairs with strongly conserved correlation values and produces a gene interaction network using a KNN algorithm.

### Network topology analysis

The topology of the network was analyzed using the R library igraph ^23,24^. The topological parameters for (i) clustering coefficient, (ii) degree assortativity, and (iii) ratio of interactions adjacent to the most significant “hub” genes (defined as 1% of genes with the highest number of interactions), were calculated compared to 1,000 randomized KNN (k=5) networks in which the five top interactions were replaced by random genes.

### Tree-and-Leaf Network

The Tree-and-Leaf (TLN) representation of the network was constructed by the following steps i-vi. (i) All gene-gene distances in the network were calculated using the Bellman-Ford method implemented in igraph. An interaction weight of 1 was assigned for all five interactions for each gene in the network, then 0.1 was added per order of interaction weight (i.e. resulting in five interactions with weights of 1 to 1.4 for each gene, with higher weights representing longer distance). (ii) The distances were structured as an average linkage hierarchical clustering dendrogram, using the R function hclust. (iii) The dendrogram was cut with an adaptive pruning algorithm to create gene clusters, using the cutreeHybrid function in the R library dynamicTreeCut. (iv) The average linkage dendrogram was then re-calculated, for the average distance between genes in each cluster above the cuts, using the R function hclust. (v) The TLN branch structure was re-created from the re-calculated dendrogram as an igraph object. Each split and branch end, represented by a node, correspond to branches and leaves in the TLN, respectively. The height of each merge in the re-calculated dendrogram was converted into an interaction weight in the TLN. (vi) The layout of the TLN was organized manually using Cytoscape. The size of each leaf node indicates the number of genes (surface area scaled according to the number of genes), the opacity of interactions indicates weights.

### Data availability

Code to perform analysis using the Liver DINA Resource will be made available upon publication. Supplementary tables are available at https://doi.org/10.5281/zenodo.15040422.

## Results

### Transcriptomic datasets included in the Liver DINA Resource

To create the Liver DINA Resource, we first manually curated a catalog of published mammalian liver datasets, publicly available from NCBI GEO ^19,20^. As the robustness of the resource is dependent upon both the quality and extent of the transcriptomic data, we used moderately strict inclusion criteria, including a minimum threshold of ≥ 5,000 genes annotated with *Homo sapiens* Ensembl gene IDs, and ≥ 10 samples in the GEO series (Fig. 1A). We also performed manual classification of the main experimental condition of each dataset, to ensure that no single condition was over-represented in our resource. The final dataset catalog includes 655 datasets, encompassing 48,311 liver transcriptomes (Supplementary Table 1).

**Fig. 1.**
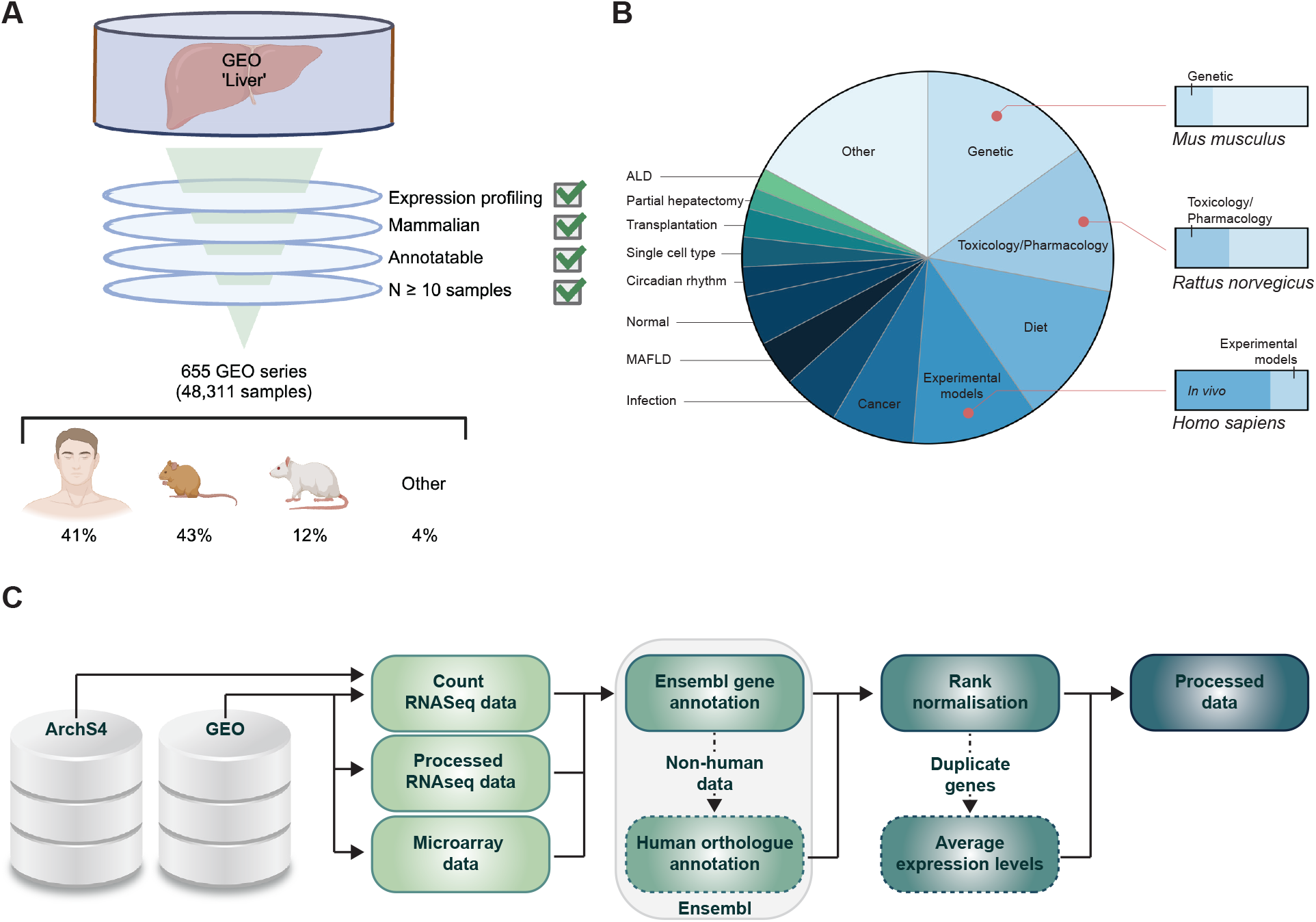
Transcriptomic datasets included in the Liver DINA Resource. **(A)** Flow chart of inclusion criteria for liver datasets. The percentage of the total sample number is shown for each species. Created in BioRender. Ankarklev, J. (2025) https://BioRender.com/r31i522. **(B)** Pie chart of classification of the experimental condition for each GEO series dataset (left), with bar charts showing the main experimental condition for each species (right). **(C)** Flow chart of the data processing pipeline.

The samples are 41% *Homo sapiens*, 43% *Mus musculus*, 12% *Rattus norvegicus*, and 4% other mammalian species (Fig. 1B). Classification of the main experimental condition of each GEO series showed that the majority of datasets are analysis of *in vivo* liver samples (89%). The main experimental conditions of the datasets include single gene knock-out or knock-down, toxicology and pharmacology treatments, diet, studies in experimental models, cancer, infection, and MAFLD, in addition to normal liver (Fig. 1B). Some experimental conditions are predominated by datasets from certain species, for example 28% of *Homo sapiens* datasets are from *in vitro* or *ex vivo* experimental models, 26% of *Mus musculus* datasets are from genetic models, while 40% of *Rattus norvegicus* datasets are from toxicology and pharmacology studies (Fig. 1B).

Our pipeline for individually processing each dataset in preparation for integration (Fig. 1C) entailed raw expression data being annotated with human Ensembl gene IDs, with data from non-human species annotated only with human one-to-one gene orthologues ^22^, and genes that did not meet a minimum threshold of unique expression values being removed from each dataset. Together, these steps ensured that the final processed datasets prioritized quality over quantity, enhancing the reliability of downstream analyses.

### Construction of the Liver DINA Resource gene-gene interaction network

As the premise of the Liver DINA Resource is centered around identifying gene-gene interactions that are preserved across mammalian liver datasets, the gene correlation structure of each of the datasets used to build the network must be consistent. To ensure this is the case, we calculated the cluster similarity of co-expressed genes between datasets and compared it to randomly produced gene clusters. Pairwise similarity between gene clustering results of datasets is significantly higher than random for 95.3% of comparisons (z-score >1.64, corresponding to *p*<0.05) (Fig. 2A). Furthermore, all datasets share significant similarity with multiple other datasets (≥24); this suggests that rather than being outliers of poor quality, datasets with unusual gene correlation structures have biological relevance that can inform the gene-gene interaction network. When the similarities between the gene clusters of the datasets are visualized as a network, datasets separate based on species. However, 16.3% of dataset-dataset interactions in the network connect datasets between species (Fig. 2B). Together, these analyses show that the similarity in gene clustering between all datasets is statistically significant, and that distinct biological factors such as species can be distinguished between the gene-gene correlation structures. Therefore, the 655 processed datasets strike a balance between the general correlation of gene expression and conditions that contribute unique information to the network.

**Fig. 2.**
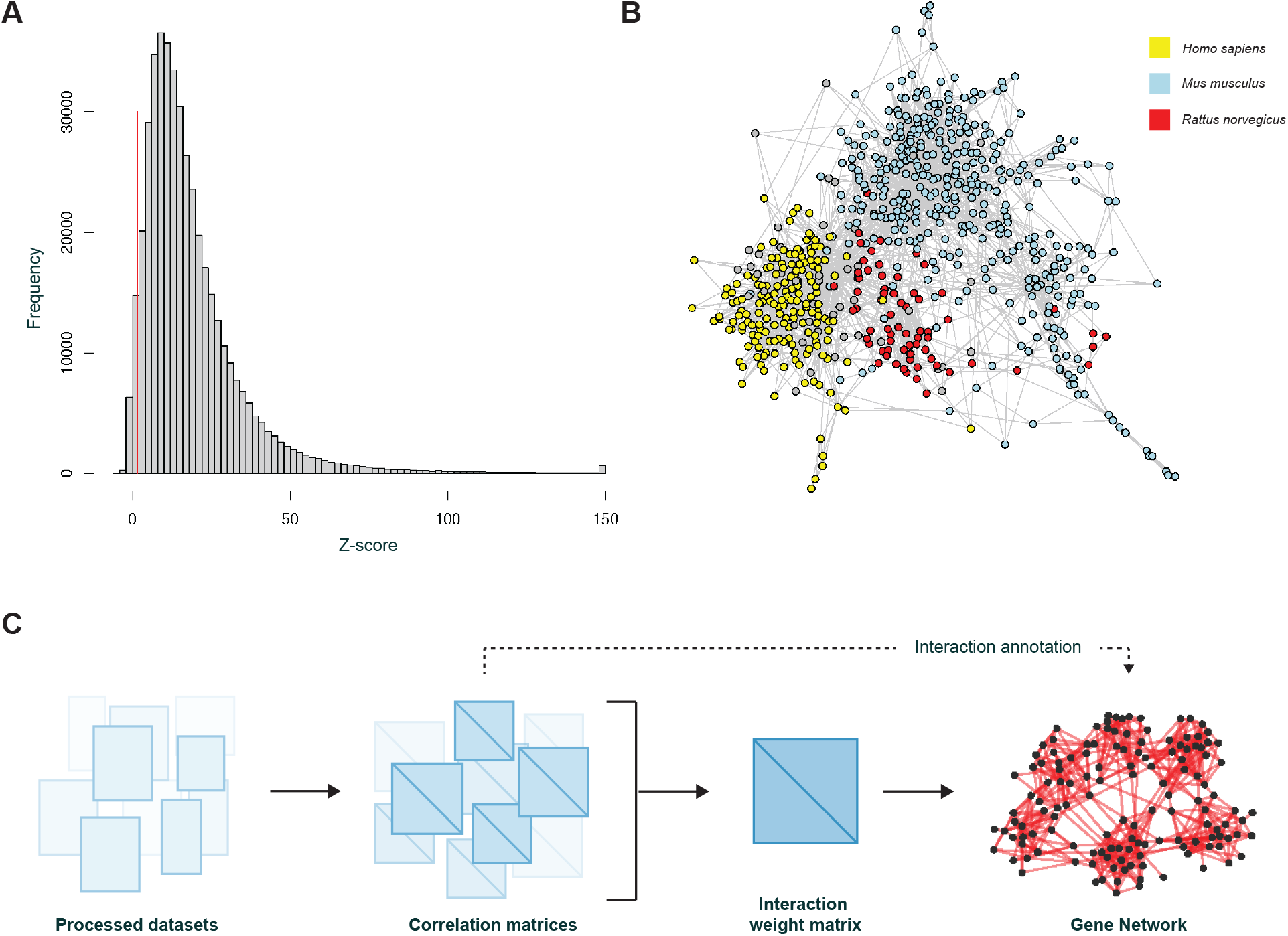
Construction of the Liver DINA Resource gene-gene interaction network. **(A)** Distribution of z-scores of clustering similarity of normalized expression of genes between datasets, compared to randomized gene clusters, measured using NMI. The threshold of significance is shown by the red line at X=1.64 (corresponding to *p*>0.05). **(B)** Network of k-nearest neighbor (KNN) similarity between gene-gene correlation structures of datasets (k=4). Yellow nodes represent *Homo sapiens* datasets, red nodes represent *Rattus norvegicus* datasets, and blue nodes represent *Mus musculus* datasets. **(C)** Liver DINA Resource network construction pipeline. Gene-gene correlation values were calculated for each processed dataset separately. All 655 correlation matrices were collapsed into a single gene-gene interaction weight matrix which was used to create a gene interaction network.

To construct a comprehensive and unbiased gene network specific to the liver, we used a pipeline that identifies statistically conserved gene-gene interactions and integrates them into a unified network for analysis (Fig. 2C). We assessed the network topology compared to 1000 randomized versions of the network. The clustering coefficient is 0.059 (95% confidence interval (CI) of random networks 3.7×10^−4^ - 5.2×10^−4^). The network has scale-free degree distribution, and the 1% of genes with highest degree (i.e. number of interactions) are collectively adjacent to 12,033 interactions (95% CI 3226-3292). Degree assortativity falls within the randomly expected range (0.043; 95% CI of random networks −0.054 − −0.042), indicating that hub genes do not preferentially interact with other hub genes. Taken together, the final network of 19,317 genes and 89,683 interactions has a modular topology with well-connected hubs, mapping the landscape of gene-gene interactions of the mammalian liver.

### Liver DINA Resource infers gene functions and pathways

As an initial assessment of the Liver DINA Resource gene-gene interaction network, we interrogated the 1,000 gene-gene interactions with the highest statistical weight. The resulting network of 588 genes retained the modularity of the full network (Fig. 3A and Supplementary Table 2). Examination of the strongest gene-gene interactions illustrates the veracity of the network. The strongest interaction - i.e. the most preserved interaction throughout a broad range of processes - connects two genes encoded by the same bicistronic transcript, a rare occurrence in mammals, *PIGY* and *PYURF* (Fig. 3B) ^25^. The second and third strongest interactions connect *C1QA, C1QB*, and *C1QC*, encoding the A- B- and C-chains of the complement component C1q ^26^. However, even the strongest interactions can also offer novel insights. Among the top 20 strongest interactions, *MANF* interacts with *CRELD2* and *SDF2L1*, all of which encode endoplasmic reticulum (ER) resident proteins involved in protein homeostasis and stress adaptation ^27–30^. Their interaction in our hepatic gene network may indicate a role in hepatic cell regulation and their potential co-expression and functional interplay in the liver warrants further investigation. Another strong interaction network involves *IDI1, MSMO1, SQLE* and *CYP51*, encoding key enzymes in cholesterol biosynthesis whose potential co-expression in the liver has not been explored ^31–33^.

**Fig. 3.**
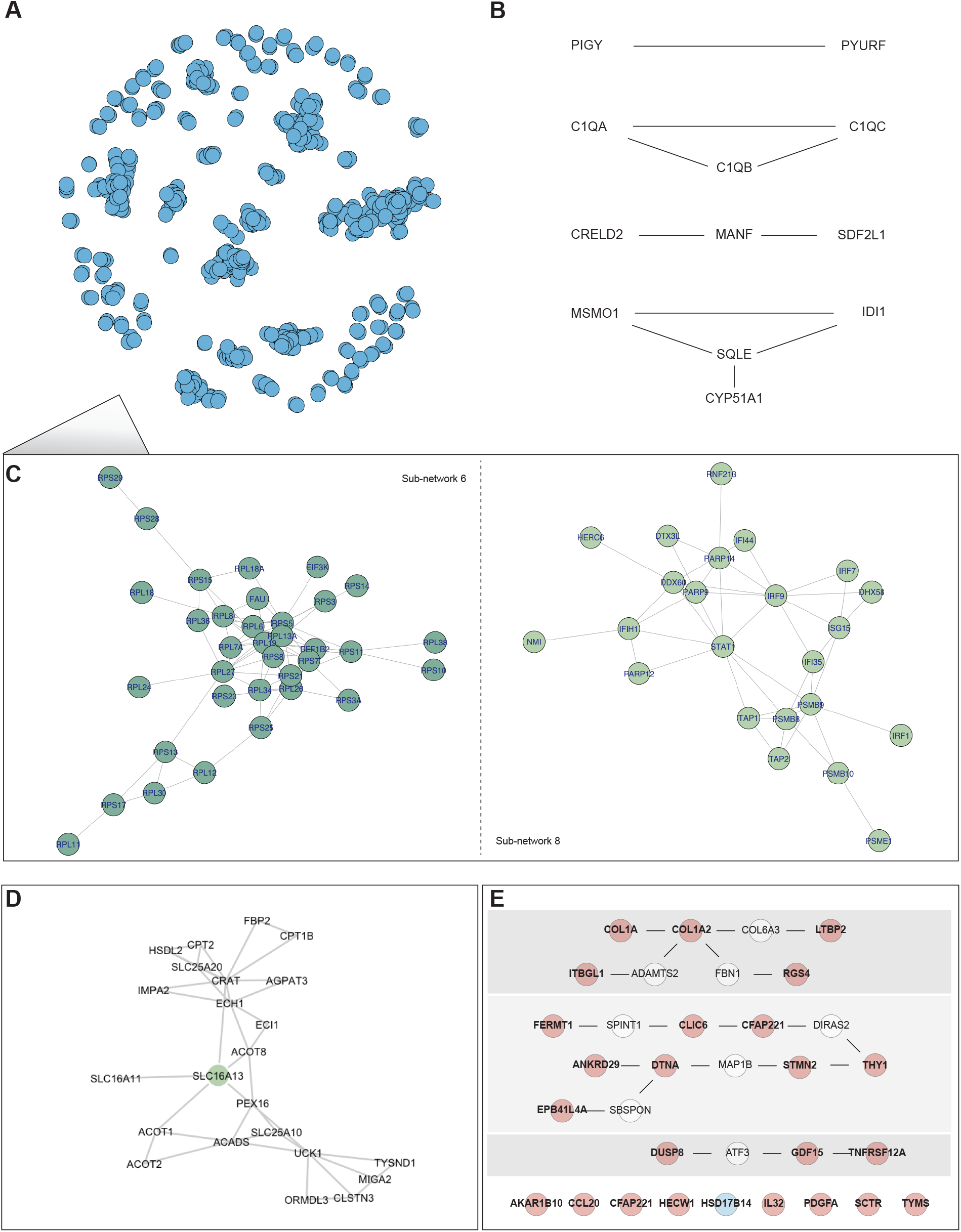
Liver DINA Resource infers gene functions and pathways. **(A)** Network diagram of the subset of gene-gene interactions with the highest statistical weight (N=1,000) in the Liver DINA Resource gene network. **(B)** Gene-gene interactions among the top 20 strongest interactions in the Liver DINA resource network. **(C)** Details of selected gene-gene interaction sub-networks within the N=1,000 subset. Sub-networks 6 and 8, represent the sixth and eighth largest network in the N=1,000 subset, respectively. **(D)** Gene interaction network for *SLC16A13* (to a maximum distance of two gene-gene interactions). **(E)** Gene networks connecting a ‘signature’ of 25 genes with a gradual increase (red) or decrease (blue) during MAFLD disease progression (with ≤1 intermediate gene-gene interaction).

The connected sub-networks formed by the top 1,000 interactions also illustrate the validity of the Liver DINA Resource, as well as its ability to reach beyond tools that depend upon our existing knowledge of genes and pathways (Supplementary Table 2 and Fig. 3C). For example, sub-network 6 (i.e. the sixth largest network) almost exclusively contains genes encoding ribosomal subunit proteins (RPs), as well as a fusion protein of RPS30 and the ubiquitin-like protein FUBI (FAU). The two additional genes in the sub-network, encoding EIF3K and EEF1B2, are less clearly connected; however, recent reports offer experimental support for their inclusion, hinting that preferential translation of RPs contributes to the modulation of global protein synthesis by these translation factors ^34,35^. In another example, sub-network 8 contains genes with firmly experimentally established interactions associated with the immune response, particularly the Type-I interferon (IFN) antiviral immune response. The sub-network is centered around the genes encoding the Type-I IFN activated transcription factors STAT1 and IRF9, and 23/23 genes in the network are Type-I IFN stimulated genes (ISGs) ^36^. Overall, the prominence of these gene-gene interactions in the Liver DINA Resource suggests these pathways are of particular interest for liver studies.

The Liver DINA Resource gene-gene interaction network can be used to infer novel gene functions or pinpoint relevant pathways within gene lists. For example, *SLC16A13*, encoding a lactate transporter, was identified as a novel susceptibility gene for type 2 diabetes ^37,38^, and *Slc16a3* deletion in a mouse model led to increased mitochondrial respiration in the liver resulting in reduced lipid accumulation and increased insulin sensitivity ^38^. The gene interaction network of *SLC16A13* in the liver suggests that it interacts with two different gene clusters, centered around *CRAT* and A*CADS*, which encode mitochondrial enzymes involved in Acyl-CoA metabolic process (Fig. 3D). In another example, interrogation of the relationship between a list of 25 genes identified through RNA sequencing analysis as a ‘signature’ associated with MALFD disease progression indicates three possible interaction pathways ^39^ (Fig. 3E). For example, one cluster involves known MAFLD markers (*COL1A1* and *COL1A2*) with *RGS4*, a gene which is known to play a role in cardiometabolic diseases but which has not been studied in the context of liver fibrosis ^40^. This network also includes *FBN1*, a gene that has been suggested by one other group to be a marker for MAFLD, but which has not been further validated ^41^. Another cluster includes *THY1*, which is an established marker of liver fibrosis and has been implicated in MAFLD progression ^42,43^. However, this network also highlights *CLIC6*, encoding a chloride intracellular channel protein not previously studied in the context of MAFLD, as well as *CFAP221* and *DIRAS2*, which have no known links to liver disease. These types of analyses allow the data-driven development of hypotheses for further investigation.

### Tree-and-Leaf Network for gene enrichment analysis of datasets

The Liver DINA Resource gene-gene interaction network is a powerful tool to explore genes and gene sets, and generate hypotheses about gene function. In addition, the modularity of the complete network can be exploited to analyze patterns in larger datasets, for example, to prioritize genes and pathways for further investigation. To facilitate such analyses, we created a streamlined visualization of the 19,317 gene network as a Tree-and-Leaf Network (TLN). Here, each gene module is represented as a leaf and their relative distance as branches (Fig. 4A). The TLN shows a highly branched topology, supporting the many diverse and adaptable functions of the liver.

**Fig. 4.**
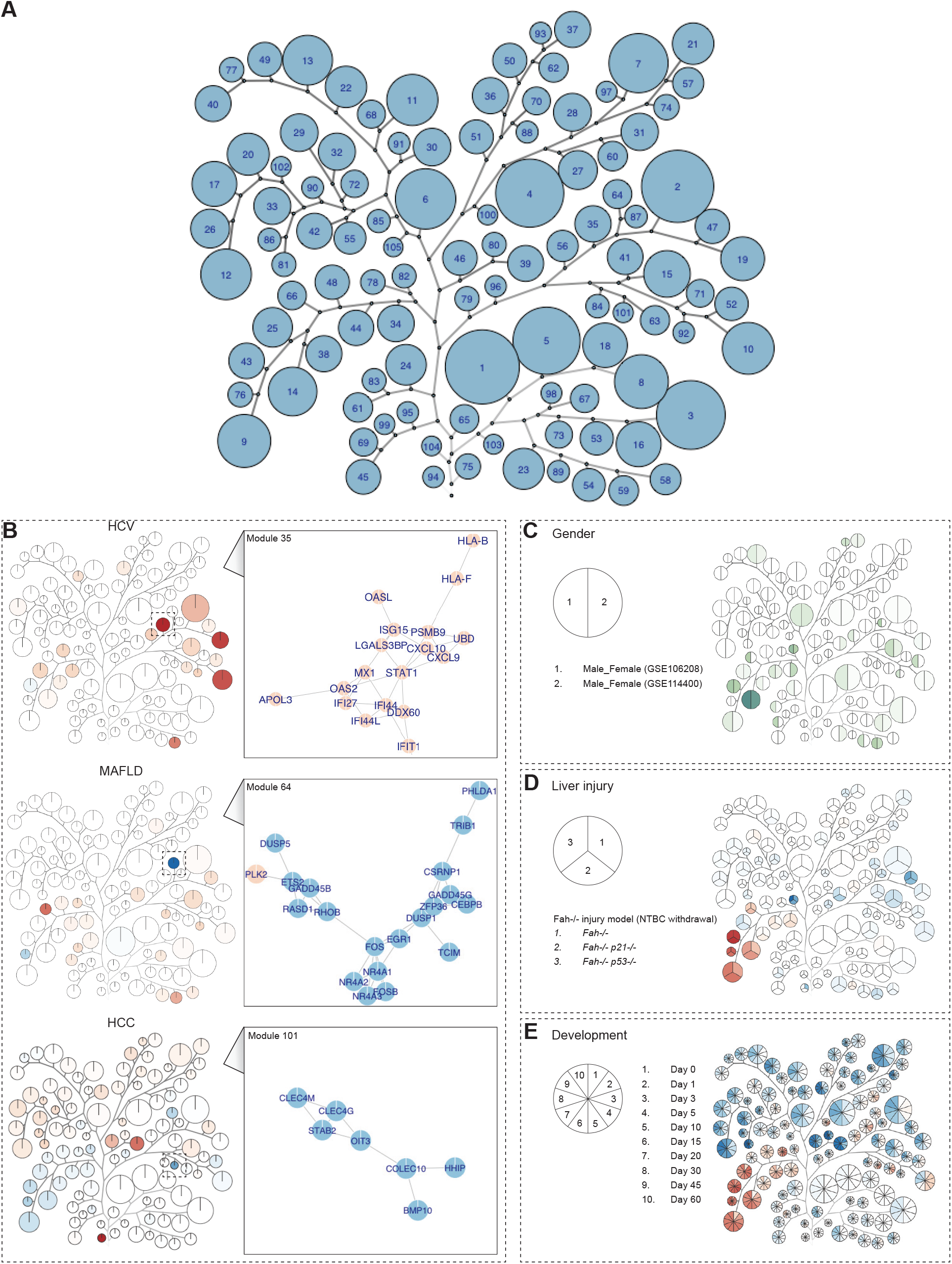
Tree-and-Leaf Network for gene enrichment analysis of datasets. **(A)** TLN visualization of the Liver DINA Resource. The size of the leaves indicates the number of genes in each cluster, with leaves labeled 1-105 in order of size (Module labels in Supplementary Fig. 1). **(B)** Visualization of changes in gene expression from publicly available expression profiling datasets. (Left) Each module is color-coded to represent the proportion of upregulated DE genes (red) to downregulated DE genes (blue), normalized to the number of genes in the module. Only genes in both (i) the dataset’s full gene list and (ii) the Liver DINA Resource are included. HCV cirrhosis compared to normal liver (DE genes adjusted *p*-value <0.0001 & logFC>log2(5)). MAFLD compared to normal liver (DE genes adjusted *p*-value <0.0001 & logFC>log2(5)). HCC compared to normal liver (DE genes adjusted *p*-value <0.05 & logFC>log2(1.5)). (Right) The largest cluster of DE genes for a specific module is shown, with upregulated genes (red) and downregulated genes (blue). (HCV: adjusted *p*-value <0.0001 & logFC>log2(2); MAFLD: adjusted *p*-value <0.0001 & logFC>log2(1.5): HCC: adjusted *p*-value <0.05 & logFC>log2(1.5)). **(C)** Comparison of DE genes in two independent datasets comparing male and female mice. For each module the magnitude of DE genes (both up and downregulated) is shown in green (DE genes adjusted *p*-value <0.0001 & logFC>log2(5)). **(D)** Changes in gene expression between *Fah*^*−/−*^ mice supplemented with NTPC (baseline), compared to *Fah*^*−/−*^, *Fah*^*−/−*^*p21*^*−/−*^, and *Fah*^*−/−*^*p53*^*−/−*^ mice following NTPC withdrawal (liver injury) (DE genes adjusted *p*-value <0.0001 & logFC>log2(5)). **(E)** DE genes during mouse development compared to late embryonic stage. The magnitude of upregulated DE genes (red) to downregulated DE genes (blue) is shown for each developmental stage (DE genes adjusted *p*-value <0.05 & logFC>log2(1.5)).

Classification of the modules using standard Gene Ontology (GO) enrichment analysis shows that some modules reflect well established gene-gene interactions (Supplementary Table 3). For example, Module 104 shows strong statistical enrichment of the Biological Process ‘Homophilic cell adhesion via plasma membrane adhesion molecules’ (FDR q-value 5.41E-90, Enrichment 108.37) and consequently is labeled as ‘Cell adhesion’. Module 45, which shows strong statistical enrichment of the Biological Process ‘Regulation of transcription by RNA polymerase II’ (FDR q-value 1.52E-81, Enrichment 5.97) is labeled as ‘Regulation of RNA Pol II transcription (I)’ (Supplementary Fig. 1). Some modules encompass several related GO Biological Processes that also reflect expected gene-gene interactions, for example, Module 46 which shows similar statistical enrichment of the Biological Process ‘Cell cycle process’, ‘DNA metabolic process’, and ‘Mitotic cell cycle process’ and consequently is labeled as ‘Cell cycle (I)’ (Supplementary Fig. 1). As expected, the branches of the TLN often connect related Biological Processes. Importantly, most modules have no clear enrichment for one specific GO Biological Process (Supplementary Table 3). These gene-gene interaction clusters are not easily explained by our current knowledge and bring less defined, potentially novel connections to the foreground. This illustrates the value of the Liver DINA Resource in uncovering relevant gene-gene interactions that would be obscured by analysis based only upon well-established gene ontologies.

Simple visualization of expression profiling datasets can help highlight genes and pathways of interest. We can project changes in gene expression onto the TLN, and identify modules enriched in differentially expressed (DE) genes. We can also visualize which genes and gene networks are DE within a module of interest. As a general example, using publicly available liver expression profiling datasets of liver disease in patients - including hepatitis C virus (HCV) (GSE14323 ^44^), MAFLD (GSE135251 ^39^), and HCC (GSE45050 ^45^) - we can observe that DE genes associate with distinctly different modules for each of the conditions (Fig. 4B). Accordingly, patients with HCV cirrhosis show enrichment of upregulated genes in the branch associated with immune response, particularly Module 35 (‘Immune response (II)’, defense response to virus). For patients with MAFLD, there is enrichment of upregulated genes in Module 66 ‘Cholesterol Biosynthesis’, which cluster around *FASN*, encoding a key enzyme in fatty acid metabolism which also promotes inflammatory immune responses ^46^ (not shown), and enrichment in downregulated genes in Module 64 (‘Regulation of cell death’), which cluster around *FOS* and *DUSP1*. Patients with HCC show enrichment of upregulated genes particularly in modules associated with hyperproliferation, Module 94 (‘Nucleosome assembly’) and Module 39 (‘Mitosis’) (not shown), as well as enrichment in downregulated genes in Module 101 (‘Tissue Morphogenesis’) which cluster around *OIT3*, typically downregulated in HCC ^47^.

The TLN can also be used to compare and contrast conditions easily. In a simple example, the DE genes of two independent datasets comparing conventionally reared male and female mice controlled for different conditions (circadian clock (GSE114400 ^48^), and mouse strain (GSE106208 ^49^) largely overlap, with Module 14 (‘Steroid Hormone Biosynthesis’) identified with high confidence. Alternatively, the contrast in response to liver injury following withdrawal of nitisinone (NTPC) in *Fah*^*−/−*^ mice can be visually examined for *Fah*^*−/−*^, *Fah*^*−/−*^*p21*^*−/−*^, and *Fah*^*−/−*^*p53*^*−/−*^ mice (GSE156852 ^50^), with Module 43 (‘Protein activation cascade’) and Module 82 (on the same branch) enriched in DE genes following liver injury. Module 82 is particularly impacted by the abrogation of p53, with downregulated expression of genes including for example *DAPK2* (Fig. 4D). In a final example, sequential changes in global gene expression states can be simply visualized using the TLN, as shown for mice developing from late embryonic stage (two days before birth) to maturity (60 days old) (GSE58827 ^51^). This shows that connected modules are typically up- or down-regulated similarly over time (Fig. 4E).

## Conclusions

The exponentially increasing public catalog of transcriptomic studies from relevant species, tissues, diseases, or other experimental conditions are a powerful asset. They enable entirely data-driven analysis for deeper insight into complex biological processes, without relying upon previous knowledge. Here, we establish Deep Integrated Network Analysis (DINA) as a valuable approach. As the liver is a widely studied organ, the Liver DINA Resource integrated a rigorous collection of liver transcriptomic datasets, including a strong foundation of *in vivo* and human studies, as well as a wide representation of common diseases and other conditions. This provides a comprehensive framework for comparative and integrative liver research across multiple experimental approaches and species. The same DINA framework can be applied to any specialized biological condition, limited only by access to appropriate datasets.

The Liver DINA Resource gene-gene interaction network, of 19,317 genes and 89,683 interactions, maintains a decentralized and scale-free topology, effectively capturing key gene-gene interactions in the mammalian liver. The network presents both well-established and novel biological interactions in the liver, providing a valuable tool for uncovering previously unrecognized functional pathways. As the network is modular, we can also visualize the higher topology as a Tree-And-Leaf Network, with ‘leaves’ of interacting genes representing many diverse functions of the liver, organized on ‘branches’ according to their relationship to each other.

Here, we have demonstrated the utility of the Liver DINA Resource as a tool for translational research, providing examples of how it can be used to explore genes and gene pathways, as well as analyze patterns in larger datasets. The interactive version of the Liver DINA Resource will be made publicly available as a platform for uncovering new biological insights in liver research.

## References

1. Devarbhavi H, Asrani SK, Arab JP, Nartey YA, Pose E, Kamath PS. Global burden of liver disease: 2023 update. J Hepatol. 2023;79(2):516–537.

2. Huang DQ, Terrault NA, Tacke F, et al. Global epidemiology of cirrhosis - aetiology, trends and predictions. Nat Rev Gastroenterol Hepatol. 2023;20(6):388–398.

3. Chidambaranathan-Reghupaty S, Fisher PB, Sarkar D. Hepatocellular carcinoma (HCC): Epidemiology, etiology and molecular classification. Adv Cancer Res. 2021;149:1–61.

4. Guilliams M, Bonnardel J, Haest B, et al. Spatial proteogenomics reveals distinct and evolutionarily conserved hepatic macrophage niches. Cell. 2022;185(2). doi:10.1016/j.cell.2021.12.018

5. Zhao X, Wang S, Liu Q, et al. Single-cell landscape of the intrahepatic ecosystem in alcohol-related liver disease. Clin Transl Med. 2025;15(1):e70198.

6. Afriat A, Zuzarte-Luís V, Bahar Halpern K, et al. A spatiotemporally resolved single-cell atlas of the Plasmodium liver stage. Nature. 2022;611(7936):563–569.

7. Hildebrandt F, Iturritza MU, Zwicker C, et al. Host-pathogen interactions in the Plasmodium-infected mouse liver at spatial and single-cell resolution. Nat Commun. 2024;15(1):7105.

8. Jiang P, Jia H, Qian X, et al. Single-cell RNA sequencing reveals the immunoregulatory roles of PegIFN-α in patients with chronic hepatitis B. Hepatology. 2024;79(1):167–182.

9. Perry AS, Hadad N, Chatterjee E, et al. A prognostic molecular signature of hepatic steatosis is spatially heterogeneous and dynamic in human liver. Cell Rep Med. 2024;5(12):101871.

10. Fred RG, Steen Pedersen J, Thompson JJ, et al. Single-cell transcriptome and cell type-specific molecular pathways of human non-alcoholic steatohepatitis. Sci Rep. 2022;12(1):13484.

11. Watson BR, Paul B, Rahman RU, et al. Spatial transcriptomics of healthy and fibrotic human liver at single-cell resolution. Nat Commun. 2025;16(1):319.

12. De Ponti FF, Bujko A, Liu Z, et al. Spatially restricted and ontogenically distinct hepatic macrophages are required for tissue repair. Immunity. 2025;58(2):362–380.e10.

13. Warde-Farley D, Donaldson SL, Comes O, et al. The GeneMANIA prediction server: biological network integration for gene prioritization and predicting gene function. Nucleic Acids Res. 2010;38(Web Server issue):W214–20.

14. Snel B, Lehmann G, Bork P, Huynen MA. STRING: a web-server to retrieve and display the repeatedly occurring neighbourhood of a gene. Nucleic Acids Res. 2000;28(18):3442–3444.

15. Szklarczyk D, Kirsch R, Koutrouli M, et al. The STRING database in 2023: protein-protein association networks and functional enrichment analyses for any sequenced genome of interest. Nucleic Acids Res. 2023;51(D1):D638–D646.

16. Buzzao D, Persson E, Guala D, Sonnhammer ELL. FunCoup 6: advancing functional association networks across species with directed links and improved user experience. Nucleic Acids Res. 2025;53(D1):D658–D671.

17. Altman MC, Rinchai D, Baldwin N, et al. Development of a fixed module repertoire for the analysis and interpretation of blood transcriptome data. Nat Commun. 2021;12(1):4385.

18. Khatri P, Sirota M, Butte AJ. Ten years of pathway analysis: current approaches and outstanding challenges. PLoS Comput Biol. 2012;8(2):e1002375.

19. Edgar R, Domrachev M, Lash AE. Gene Expression Omnibus: NCBI gene expression and hybridization array data repository. Nucleic Acids Res. 2002;30(1):207–210.

20. Barrett T, Wilhite SE, Ledoux P, et al. NCBI GEO: archive for functional genomics data sets--update. Nucleic Acids Res. 2013;41(Database issue):D991–5.

21. Lachmann A, Torre D, Keenan AB, et al. Massive mining of publicly available RNA-seq data from human and mouse. Nat Commun. 2018;9(1):1366.

22. Dyer SC, Austine-Orimoloye O, Azov AG, et al. Ensembl 2025. Nucleic Acids Res. 2025;53(D1):D948–D957.

23. Csardi G, Nepusz T. The igraph software package for complex network research. InterJournal, Complex Systems. 2006;1695.

24. Csárdi G, Nepusz T, Traag V, et al. igraph: Network Analysis and Visualization in R. 2025. doi:10.5281/zenodo.7682609 <https://doi.org/10.5281/zenodo.7682609>, R package version 2.1.4.9014, <https://CRAN.R-project.org/package=igraph>

25. Murakami Y, Siripanyaphinyo U, Hong Y, Tashima Y, Maeda Y, Kinoshita T. The initial enzyme for glycosylphosphatidylinositol biosynthesis requires PIG-Y, a seventh component. Mol Biol Cell. 2005;16(11):5236–5246.

26. Reid KBM. Complement component C1q: Historical perspective of a functionally versatile, and structurally unusual, serum protein. Front Immunol. 2018;9:764.

27. Apostolou A, Shen Y, Liang Y, Luo J, Fang S. Armet, a UPR-upregulated protein, inhibits cell proliferation and ER stress-induced cell death. Exp Cell Res. 2008;314(13):2454–2467.

28. Tang Q, Li Y, He J. MANF: an emerging therapeutic target for metabolic diseases. Trends Endocrinol Metab. 2022;33(4):236–246.

29. Kern P, Balzer NR, Blank N, et al. Creld2 function during unfolded protein response is essential for liver metabolism homeostasis. FASEB J. 2021;35(10):e21939.

30. Tang Q, Liu Q, Li Y, Mo L, He J. CRELD2, endoplasmic reticulum stress, and human diseases. Front Endocrinol (Lausanne). 2023;14:1117414.

31. Arbo MD, Melega S, Stöber R, et al. Hepatotoxicity of piperazine designer drugs: up-regulation of key enzymes of cholesterol and lipid biosynthesis. Arch Toxicol. 2016;90(12):3045–3060.

32. Chen W, Xu J, Wu Y, et al. The potential role and mechanism of circRNA/miRNA axis in cholesterol synthesis. Int J Biol Sci. 2023;19(9):2879–2896.

33. Kondo A, Yamamoto S, Nakaki R, et al. Extracellular acidic pH activates the sterol regulatory element-binding protein 2 to promote tumor progression. Cell Rep. 2017;18(9):2228–2242.

34. Gerashchenko MV, Nesterchuk MV, Smekalova EM, et al. Translation elongation factor 2 depletion by siRNA in mouse liver leads to mTOR-independent translational upregulation of ribosomal protein genes. Sci Rep. 2020;10(1):15473.

35. Duan H, Zhang S, Zarai Y, et al. eIF3 mRNA selectivity profiling reveals eIF3k as a cancer-relevant regulator of ribosome content. EMBO J. 2023;42(12):e112362.

36. Rusinova I, Forster S, Yu S, et al. Interferome v2.0: an updated database of annotated interferon-regulated genes. Nucleic Acids Res. 2013;41(Database issue):D1040–6.

37. SIGMA Type 2 Diabetes Consortium, Williams AL, Jacobs SBR, et al. Sequence variants in SLC16A11 are a common risk factor for type 2 diabetes in Mexico. Nature. 2014;506(7486):97–101.

38. Schumann T, König J, von Loeffelholz C, et al. Deletion of the diabetes candidate gene Slc16a13 in mice attenuates diet-induced ectopic lipid accumulation and insulin resistance. Commun Biol. 2021;4(1):826.

39. Govaere O, Cockell S, Tiniakos D, et al. Transcriptomic profiling across the nonalcoholic fatty liver disease spectrum reveals gene signatures for steatohepatitis and fibrosis. Sci Transl Med. 2020;12(572):eaba4448.

40. McNeill SM, Zhao P. The roles of RGS proteins in cardiometabolic disease. Br J Pharmacol. 2024;181(14):2319–2337.

41. Lou Y, Tian GY, Song Y, et al. Characterization of transcriptional modules related to fibrosing-NAFLD progression. Sci Rep. 2017;7(1):4748.

42. Gao R, Wang J, He X, et al. Comprehensive analysis of endoplasmic reticulum-related and secretome gene expression profiles in the progression of non-alcoholic fatty liver disease. Front Endocrinol (Lausanne). 2022;13:967016.

43. Blank V, Karlas T, Anderegg U, et al. Thy-1 restricts steatosis and liver fibrosis in steatotic liver disease. Liver Int. 2024;44(8):2075–2090.

44. Mas VR, Maluf DG, Archer KJ, et al. Genes involved in viral carcinogenesis and tumor initiation in hepatitis C virus-induced hepatocellular carcinoma. Mol Med. 2009;15(3-4):85–94.

45. Darpolor MM, Basu SS, Worth A, et al. The aspartate metabolism pathway is differentiable in human hepatocellular carcinoma: transcriptomics and (13) C-isotope based metabolomics. NMR Biomed. 2014;27(4):381–389.

46. Xiao Y, Yang Y, Xiong H, Dong G. The implications of FASN in immune cell biology and related diseases. Cell Death Dis. 2024;15(1):88.

47. Xu ZG, D. Jj, Zhang X, et al. A novel liver-specific zona pellucida domain containing protein that is expressed rarely in hepatocellular carcinoma. Hepatology. 2003;38(3):735–744.

48. Weger BD, Gobet C, Yeung J, et al. The mouse microbiome is required for sex-specific diurnal rhythms of gene expression and metabolism. Cell Metab. 2019;29(2):362–382.e8.

49. Duncan CG, Grimm SA, Morgan DL, et al. Dosage compensation and DNA methylation landscape of the X chromosome in mouse liver. Sci Rep. 2018;8(1):10138.

50. Buitrago-Molina LE, Marhenke S, Becker D, et al. P53-independent induction of p21 fails to control regeneration and hepatocarcinogenesis in a Murine liver injury model. Cell Mol Gastroenterol Hepatol. 2021;11(5):1387–1404.

51. Renaud HJ, Cui YJ, Lu H, Zhong XB, Klaassen CD. Ontogeny of hepatic energy metabolism genes in mice as revealed by RNA-sequencing. PLoS One. 2014;9(8):e104560.

52. Eden E, Lipson D, Yogev S, Yakhini Z. Discovering motifs in ranked lists of DNA sequences. PLoS Comput Biol. 2007;3(3):e39.

53. Eden E, Navon R, Steinfeld I, Lipson D, Yakhini Z. GOrilla: a tool for discovery and visualization of enriched GO terms in ranked gene lists. BMC Bioinformatics. 2009;10(1):48.

