## Supplementary material for "Deep Integrated Network Analysis – a data-driven tool to discover and characterize disease pathways in the liver"

### Supplementary Methods

*Classification of TLN modules.*

Labeling of branches and/or leaves of the Liver DINA Resource TLN was performed manually. A search of enriched GO terms in the gene list of modules compared to the background list of all genes in the Liver DINA network was performed using GOrilla [52,53]. The most significantly enriched GO_Process for each Module was listed, as well as additional GO_Process where the FDR q-value is within E-06 of the most significant value (up to a maximum of 20), only for FDR q-value < E-03.

**Supplementary Tables**

Supplementary tables are available at *https://doi.org/10.5281/zenodo.15040422.*

**Supplementary Table 1. List of datasets included in Liver DINA Resource.** For each dataset the GEO series, title, taxonomy, and liver sample count are shown, as well as the classification of dataset condition.

**Supplementary Table 2. Liver DINA Resource gene-gene interaction network subset (N=1,000 by highest statistical weight).** The N=1,000 subset has 1,000 interactions between 588 genes. The size of the gene networks is shown for all gene-gene interactions (76 networks). The gene lists are provided for all gene-gene interaction networks of larger than average size (≥10 genes). The most significantly enriched GO_Process for each of the largest networks is listed, including additional GO where the FDR q-value is within E-06 of the most significant value, up to a maximum of 20 (FDR q-value < E-03 only).

**Supplementary Table 3. Liver DINA Resource Tree-and-Leaf Network (TLN) leaf classification.** Each leaf in the TLN is designated with a Module number in order of size (according to the number of genes in the Module). The gene list is provided for each Module. The most significantly enriched GO_Process for each Module is listed, including additional GO_Process where the FDR q-value is within E-06 of the most significant value, up to a maximum of 20 (FDR q-value < E-03 only).

**Supplementary Figure 1. Gene Ontology analysis of the Liver DINA Resource Tree-and-Leaf Network (TLN).** GO ‘Biological Process’ classification of TLN leaves. Where no GO ‘Biological Process’ is significantly enriched (FDR *q*-value < E-03) for genes within the module, the leaf is unlabeled.

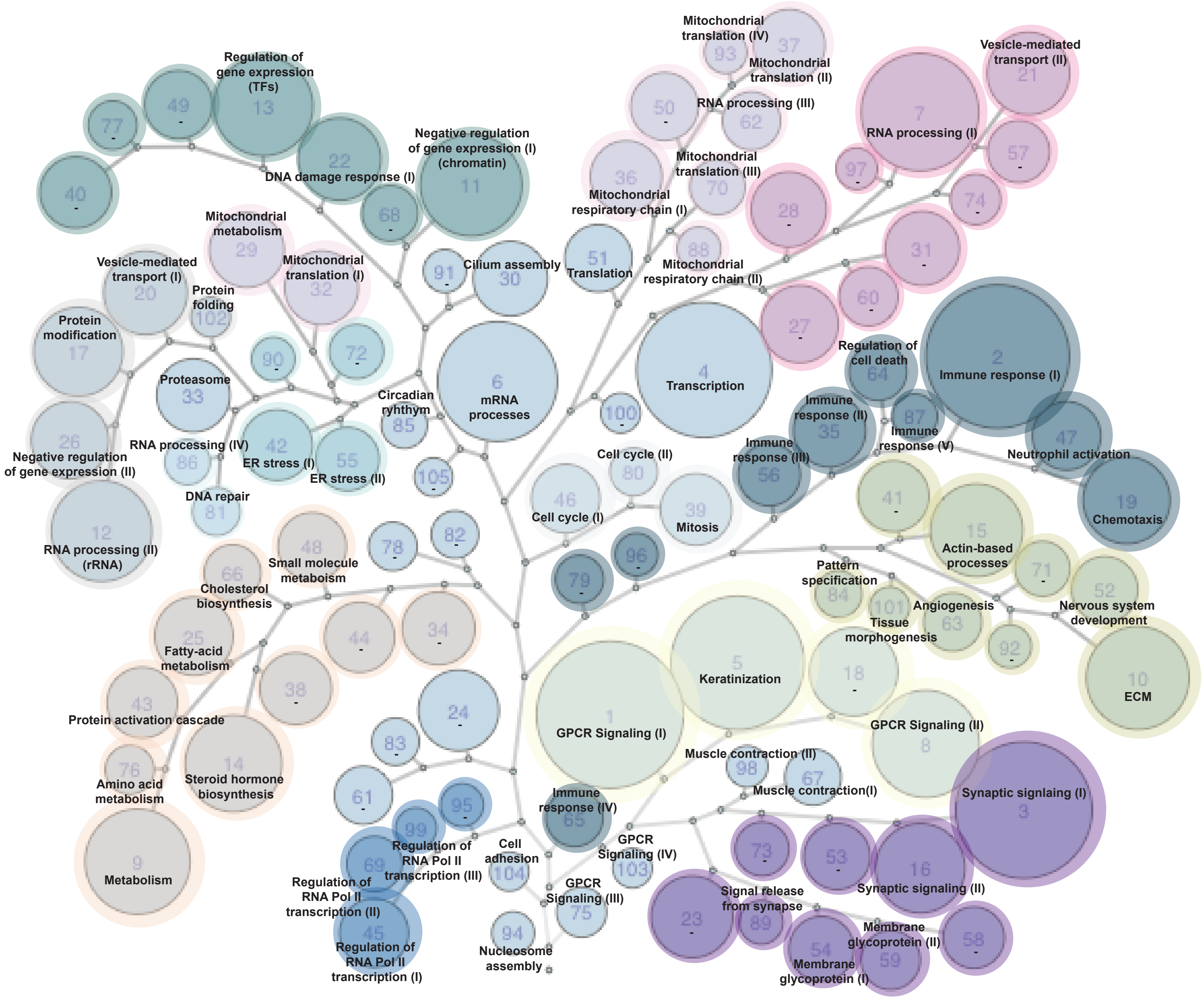
Regulation of gene expression (TFs)
DNA damage response (I)
Negative regulation of gene expression (I) (chromatin)
Mitochondrial metabolism
Vesicle-mediated transport (I)
Mitochondrial translation (I)
Protein folding
Protein modification
Cilium assembly
Translation
Proteasome
Circadian rhythym
mRNA processes
RNA processing (IV)
ER stress (I)
ER stress (II)
Negative regulation of gene expression (II)
DNA repair
RNA processing (II) (rRNA)
Small molecule metaboism
Cholesterol biosynthesis
Fatty-acid metabolism
Protein activation cascade
Amino acid metabolism
Steroid hormone biosynthesis
Metabolism
Mitochondrial translation (IV)
Mitochondrial translation (II)
RNA processing (III)
Mitochondrial translation (III)
Mitochondrial respiratory chain (I)
Mitochondrial respiratory chain (II)
Vesicle-mediated transport (II)
RNA processing (I)
Transcription
Regulation of cell death
Immune response (I)
Immune response (II)
Immune response (V)
Immune response (III)
Neutrophil activation
Chemotaxis
Cell cycle (II)
Cell cycle (I)
Mitosis
Actin-based processes
Pattern specification
Angiogenesis
Tissue morphogenesis
Nervous system development
ECM
Keratinization
GPCR Signaling (I)
GPCR Signaling (II)
Muscle contraction (II)
Muscle contraction(I)
Synaptic signlaing (I)
Immune response (IV)
Cell adhesion
GPCR Signaling (IV)
GPCR Signaling (III)
Signal release from synapse
Synaptic signaling (II)
Membrane glycoprotein (II)
Membrane glycoprotein (I)
Regulation of RNA Pol II transcription (III)
Regulation of RNA Pol II transcription (II)
Regulation of RNA Pol II transcription (I)
Nucleosome assembly
